# Mitochondrial Contact Site and Cristae Organizing System (MICOS) Machinery Supports Heme Biosynthesis by Enabling Optimal Performance of Ferrochelatase

**DOI:** 10.1101/2021.06.01.446600

**Authors:** Jonathan V. Dietz, Mathilda M. Willoughby, Robert B. Piel, Teresa A. Ross, Iryna Bohovych, Hannah G. Addis, Jennifer L. Fox, William N. Lanzilotta, Harry A. Dailey, James A. Wohlschlegel, Amit R. Reddi, Amy E. Medlock, Oleh Khalimonchuk

## Abstract

Heme is an essential cofactor required for a plethora of cellular processes in eukaryotes. In metazoans the heme biosynthetic pathway is typically partitioned between the cytosol and mitochondria, with the first and final steps taking place in the mitochondrion. The pathway has been extensively studied, and all the biosynthetic enzymes have been structurally characterized to varying extents. Nevertheless, our understanding of the regulation of heme synthesis and factors that influence this process in metazoans remains incomplete. Herein we investigate the molecular organization as well as the catalytic and structural features of the terminal pathway enzyme, ferrochelatase (Hem15), in the yeast *Saccharomyces cerevisiae*. Biochemical and genetic analyses reveal dynamic association of Hem15 with Mic60, a core component of the mitochondrial contact site and cristae organizing system (MICOS). Loss of MICOS negatively impacts Hem15 activity and results in accumulation of highly reactive and potentially toxic tetrapyrrole precursors that may result in oxidative damage. Restoring intermembrane connectivity in MICOS-deficient cells mitigates these cytotoxic effects. Our data provide new insights into how heme biosynthetic machinery is organized and regulated, linking mitochondrial architecture-organizing factors to heme homeostasis.

## INTRODUCTION

Heme is an essential cofactor and signaling molecule required for diverse physiological processes (1–4). Most metazoans synthesize heme through a highly conserved and well-characterized eight-step pathway (5). In the first step the mitochondrial enzyme aminolevulinic acid synthase (Alas; Hem1 in yeast) catalyzes the condensation of glycine and succinyl-CoA to form aminolevulinic acid (ALA), which is then transported out of the mitochondrion by an as-yet-uncharacterized transporter. Once in the cytosol, two ALA molecules are condensed into a monopyrrole, porphobilinogen, by the enzyme porphobilinogen synthase (Pbgs); subsequently, four porphobilinogen molecules are joined together by the enzyme hydroxymethylbilane synthase (Hmbs) to form the linear tetrapyrrole hydroxymethylbilane (HMB). HMB then undergoes cyclization to form uroporphyrinogen III (catalyzed by uroporphyrinogen synthase, Uros) and decarboxylation of its pyrrole acetate side chains (catalyzed by uroporphyrinogen decarboxylase, Urod) to yield coproporphyrinogen III. In *Saccharomyces cerevisiae*, coproporphyrinogen III is converted to protoporphyrinogen IX by the enzyme coproporphyrinogen oxidase (Cpox) in the cytosol (6, 7), while in humans this reaction occurs in the mitochondrial intermembrane space (IMS) (8, 9). Protoporphyrinogen IX is then transported into the mitochondrial matrix and converted to protoporphyrin IX (PPIX) by the matrix-localized enzyme protoporphyrinogen oxidase (Ppox) (10). How protoporphyrinogen IX is transported across the inner mitochondrial membrane (IM) is currently unclear though evidence suggests that the porphyrinogen transporter TMEM14C facilitates this process during erythroid cell maturation (11). The final step of heme synthesis is catalyzed by ferrochelatase (Fech; Hem15 in yeast), which resides on the matrix side of the IM (12) and mediates the insertion of ferrous iron into PPIX, thereby yielding protoheme, also known as heme *b*. Despite the fact that all the proteins involved in the synthesis of heme *b* have been structurally characterized and for some detailed kinetic studies have revealed their mechanisms of action (3, 13), the processes by which the relative rates of heme synthesis are regulated remain obscure.

Understanding of the regulation of heme synthesis is mainly limited to the transcriptional level in multicellular organisms with erythrocytes. In these organisms, most heme production occurs in the developing erythron, and specific transcription factors are known to play a role in regulating expression of all the biosynthetic enzymes (14). The mechanisms of regulation of heme synthesis for many other cells in these organisms or in unicellular eukaryotes like *S. cerevisiae*, all of which require less heme, are still uncharacterized. In terms of enzyme activity, the rate-limiting step of the pathway in mammals is the first step (catalyzed by ALAS), with the last step (catalyzed by FECH) being the second regulatory point of the mammalian pathway (14). In *S. cerevisiae*, two other pathway enzymes, Pbgs and Hmbs, have been proposed to catalyze the rate-limiting steps (7, 15). Recent data suggest additional levels of regulation at the post-translational level (16, 17).

Proteomic studies in mammalian cells provide evidence that the mitochondrial heme biosynthetic machinery exists as a large supercomplex, termed the heme metabolon (17–19). FECH was identified as a component of a multimeric assembly that includes ALAS and PPOX (17). Additional candidate interaction partners of FECH include factors that mediate IM dynamics and ultrastructure (17, 19), most notably components of the conserved mitochondrial contact site and cristae organizing system (MICOS). As suggested by its name, MICOS maintains mitochondrial cristae and contact sites between the IM and the outer mitochondrial membrane (OM) (20–22) and has been postulated to facilitate bidirectional transport of hydrophobic molecules such as phosphatidic acid (23) and coenzyme Q biosynthetic intermediates (24). However, the physiological significance of its connection to FECH remains unaddressed.

Using a yeast genetic model, we have investigated the putative molecular interaction of Fech (Hem15 in yeast) with MICOS. Data from biochemical and genetic analyses are consistent with a dynamic association of Hem15 with the core MICOS component Mic60. Loss of MICOS negatively impacts Hem15 activity and results in accumulation of cytotoxic pathway intermediates. These data provide insights into how the heme biosynthetic machinery is organized and supported, linking mitochondrial architecture to heme homeostasis.

## RESULTS

### Functional Hem15 Forms a High-Mass Complex

Hem15 exists as a homodimer (25, 26), but its protein-protein interaction has not been systematically studied. To that end, an antibody was developed against Hem15, which efficiently and specifically recognizes *S. cerevisiae* Hem15 (Fig. 1A). Using purified Hem15 as a reference, mitochondria from galactose-cultured wild type (WT) cells in the mid-log growth phase were found to contain approximately 1.0 ng of endogenous Hem15 per mg of mitochondrial protein (Supplementary Fig. S1A). This concentration is comparable to other mitochondrial heme synthesis enzymes as well as other IM proteins (27). To assess the oligomerization properties of Hem15, WT cells were fractionated, mitochondria lysed with digitonin, and the clarified lysates subjected to sucrose gradient ultracentrifugation. Hem15 was detected by SDS-PAGE immunoblot from fractions in the molecular weight range of ∼250-440 kDa (Supplementary Fig. S1B). Further investigation of this finding by the alternative technique of native gel electrophoresis was attempted, but no signal from the anti-Hem15 antibody was detected under those conditions. These experiments suggest that endogenous yeast Hem15 exists as a high-mass complex, which likely includes additional associated components, as seen for the mammalian enzyme (17).

**Figure 1.**
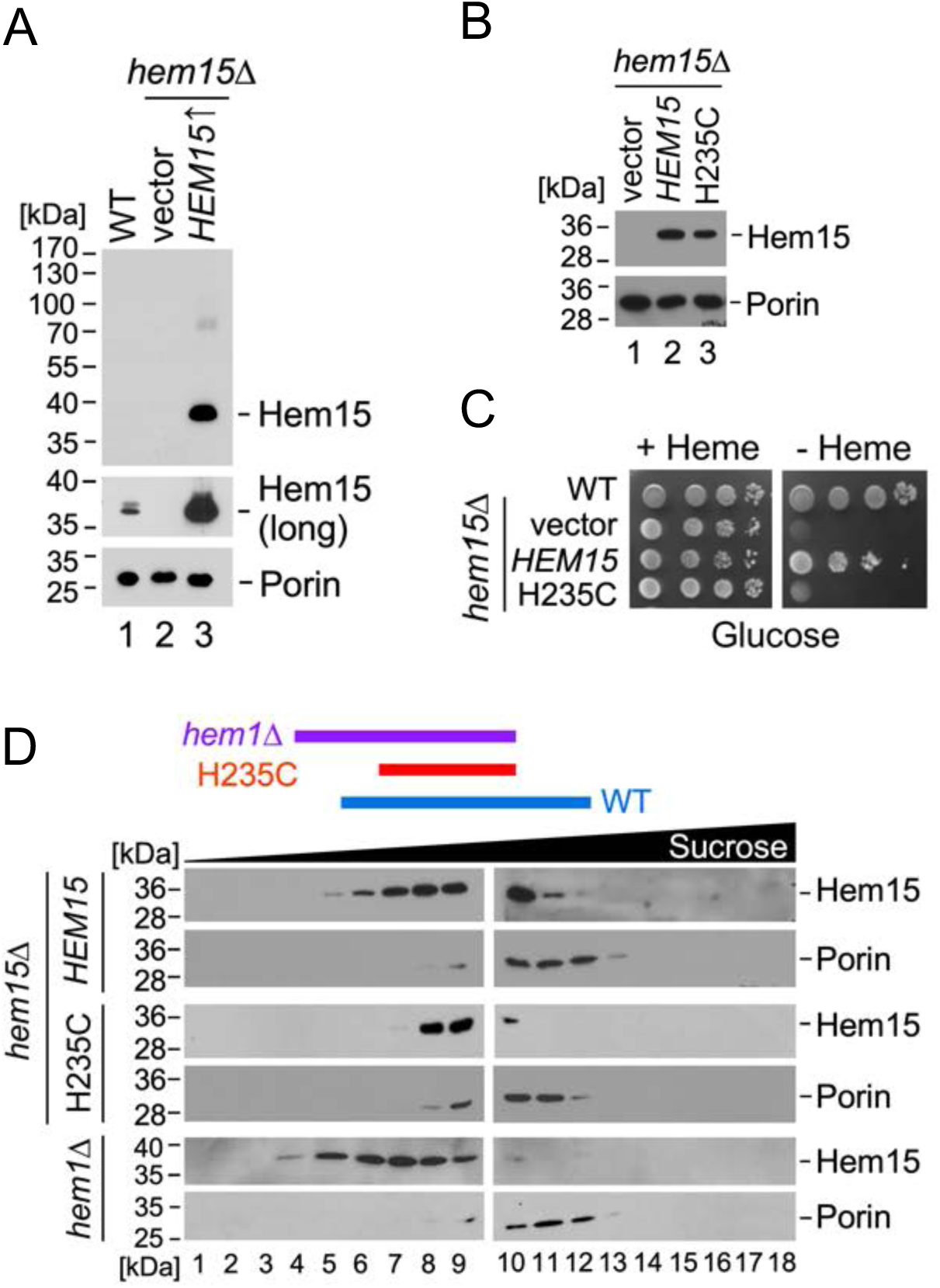
Oligomeric state of Hem15 depends on catalytic activity. **(A)** Isolated mitochondria (15 μg total protein) from wild type (WT) and *hem15*Δ cells overexpressing Hem15 (from the *HEM15* promoter) or expressing a vector control were subjected to SDS-PAGE and analyzed by immunoblotting with antibodies against Hem15 and the outer mitochondrial membrane protein Porin (loading control). **(B)** Steady-state levels of Hem15 and loading control (Porin) in mitochondria from cells described in panel A and the Hem15 variant H235C, analyzed by SDS-PAGE and immunoblotting with respective antibodies. **(C)** Heme-dependence growth test of WT cells and *hem15*Δ cells expressing vector control, Hem15 (untagged, from its native promoter on a YEp plasmid), or its H235C variant (a catalytic mutant). Cells were spotted onto SC medium with or without 20 μM hemin and cultured for 2 days at 28°C. **(D)** High-velocity density-gradient fractionation of digitonin-solubilized lysates from mitochondria of *hem1*Δ cells or *hem15*Δ cells expressing Hem15 or its H235C catalytically impaired variant, analyzed by SDS-PAGE with immunoblotting for Hem15 and Porin (440-kDa molecular size reference).

Next, the question of whether the functional state of Hem15 is important for complex formation was addressed by employing the catalytically impaired Hem15 variant H235C. The yeast Hem15 residue H235 is equivalent to the human FECH residue H263, which is an essential catalytic histidine involved in proton abstraction prior to chelation (28). When expressed in the *hem15*Δ deletion strain, the H235C variant is stable (Fig. 1B) but unable to support heme-independent growth (Fig. 1C). Unlike WT Hem15, the H235C variant does not rescue growth of the *Escherichia coli ppfC*Δ mutant and shows no detectable *in vitro* Fech activity (Supplementary Fig. S1C). When the WT and H235C variant proteins were purified and analyzed by UV-Vis spectroscopy, the WT Hem15 predominantly co-purified with bound heme (Soret *λ*_max_ = 427 nm), but the H235C variant co-purified with bound PPIX (Soret *λ*_max_ = ∼410 nm) (Supplementary Fig. S1D). To determine how the active-site structure of Hem15 is impacted by the H235C mutation, the H235C variant was crystallized and its structure solved at 2.4 Å resolution. This structure reveals that the location and orientation of active-site residues in the WT protein and H235C Hem15 variant are similar. Notably, the active-site hydrogen-bonding network of the H235C variant is intact and similar to that of the WT enzyme (Supplementary Fig. S1E), unlike the human H263C variant structure where this network is disrupted (28). It is unclear why this difference between the yeast and human catalytic mutants exists, though both variants are inactive due to the absence of the base (H235 in yeast and H263 in humans) necessary for proton abstraction.

To examine the level of oligomerization of Hem15 in the absence of enzyme function, the sucrose gradient migration profiles of the H235C Hem15 variant and the WT enzyme were compared. The H235C variant exhibited an altered migration pattern indicative of a shift toward smaller complex size, likely stemming from changes in the enzyme’s interaction with protein partners or oligomerization state (Fig. 1D). Notably, a similar size shift was observed for Hem15 in mitochondrial lysates from heme synthesis-deficient cells lacking Alas (yeast Hem1; Fig. 1D). These findings suggest that Hem15 oligomerization is at least partially dependent on the functional state of the enzyme.

### Genetic Model to Assess Hem15 Functional Roles Enables Complementation of the Yeast Deletion Mutant by Human FECH

Functional studies of Hem15 are limited due to the conditionally essential nature of the enzyme. While growth of the *hem15*Δ mutant in the presence of glucose can be sustained with either exogenous heme supplementation or re-expression of plasmid-borne Hem15 (7), this is not always the case for respiratory growth (Supplementary Fig. S2A and (29)). Since a fully functional genetic model is critical for further studies, this issue was examined in greater detail. First the heme-containing electron transport chain complexes III and IV were evaluated in mitochondria isolated from *hem15*Δ cells expressing an empty vector or YEp-*HEM15* (either under the control of its native promoter or the high-efficiency heterologous *MET25* promoter). These respiratory complexes, as well as the non-heme complex V, were severely attenuated in *hem15*Δ cells regardless of Hem15 re-expression (Fig. 2A, lanes 2, 4, and 6). Additionally, the steady-state levels of core complex IV subunits were notably reduced relative to WT control, and Hem15 re-expression did not result in complete stabilization of these subunits (Fig. 2B). Both results are consistent with the respiratory-deficient phenotype of these cells even upon Hem15 re-expression (Fig. 2C and Supplementary Fig. S2A). Because of this inability of plasmid-borne Hem15 to fully complement the *hem15*Δ mutant in some, but not all, colonies of transformed cells, we hypothesized that such an effect could stem from a petite-inducing phenotype of Hem15 deletion. To test this postulate, the dNTP checkpoint enzyme Rnr1 (encoding the large subunit of ribonucleotide reductase) was overexpressed in *hem15*Δ cells, since this enzyme has been previously shown to efficiently stabilize mitochondrial DNA in various petite mutants via an unknown mechanism (30–32). Indeed, combined expression of *RNR1* with either of the YEp-*HEM15* plasmids permits respiratory growth of the *hem15*Δ mutant (Fig. 2C) and results in marked stabilization of the mitochondrial respiratory complexes III-IV and core complex IV subunits (Fig. 2A lanes 5 and 7 and Fig. 2D lanes 5 and 7).

**Figure 2.**
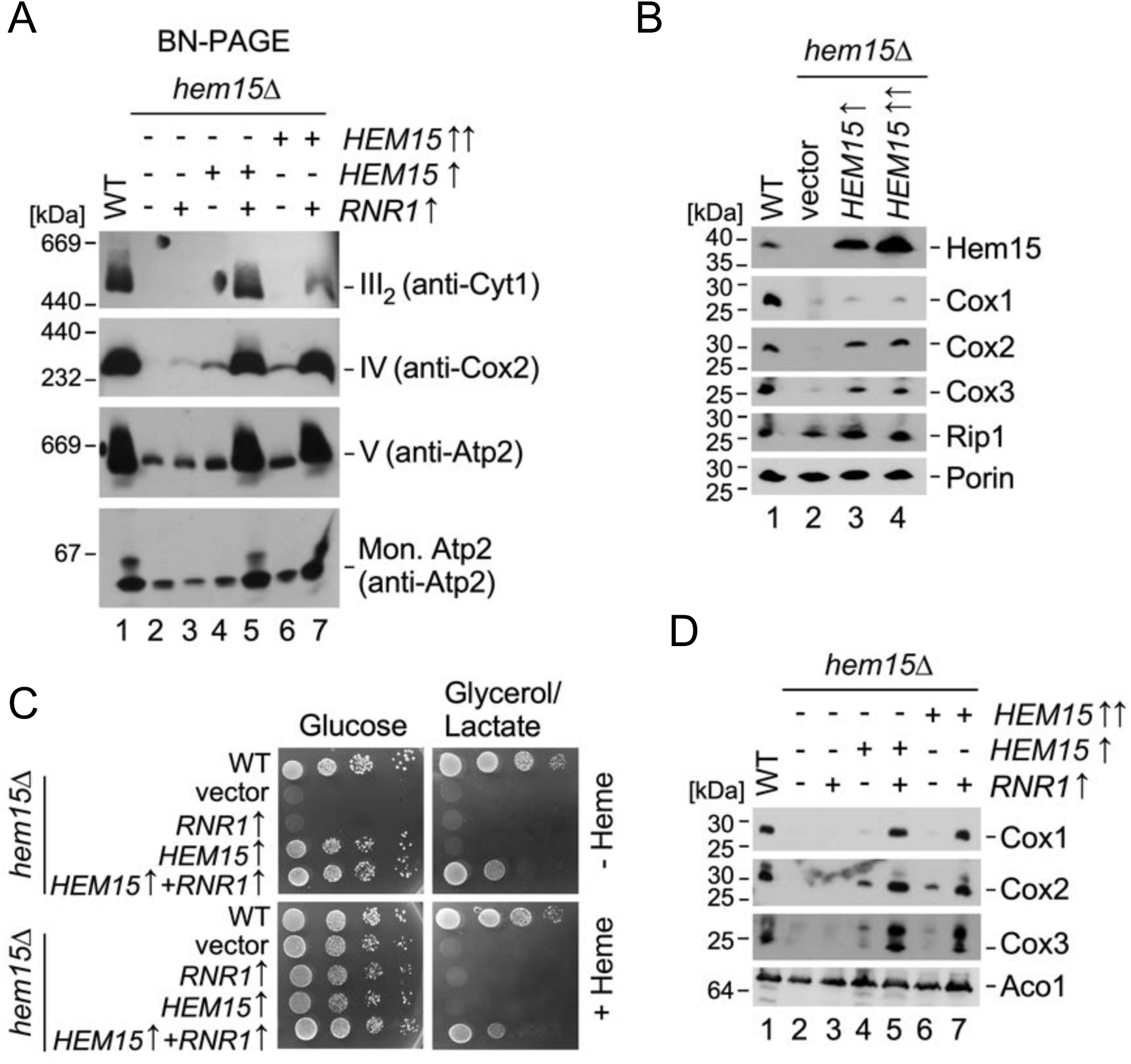
Loss of Hem15 results in a petite-inducing phenotype that can be reversed by overexpression of ribonucleotide reductase, Rnr1. **(A)** BN-PAGE analysis of dodecyl-β-D-maltoside (DDM)-solubilized mitochondrial proteins from WT cells or *hem15*Δ cells co-expressing the indicated combinations of the following plasmids: vector control (-), Rnr1 (*RNR1* ↑), Hem15 under control of its native promoter (*HEM15* ↑), and Hem15 under the control of the heterologous *MET25* promoter (*HEM15* ↑↑). Complexes were visualized by immunoblotting with the indicated antibodies (mon. Atp2 refers to the monomeric protein). **(B)** Steady-state levels of Hem15 and representative respiratory complex IV (Cox1-Cox3) and complex III (Rip1) subunits with loading control (Porin) in mitochondria of WT cells and *hem15*Δ cells with or without *HEM15* overexpression. Samples were analyzed by SDS-PAGE and immunoblotting with indicated antibodies. **(C)** Heme-dependent fermentative and respiratory growth test of cells described in panel A. Cells were spotted onto SC media with or without 20 μM hemin and cultured for 2 days (glucose) or 5 days (glycerol/lactate) at 28°C. **(D)** SDS-PAGE immunoblot showing steady-state levels of indicated mitochondrial proteins in cells described in panel A.

Interestingly, Rnr1 expression also permits complementation of the yeast *hem15*Δ mutant with human FECH, indicating that despite certain differences the human enzyme is fully functional in yeast (Supplementary Fig. S2B). This finding reveals that essential protein-protein interactions between FECH and other components are conserved in *S. cerevisiae*. The human enzyme, but not the yeast enzyme, possesses a [2Fe-2S] cluster, which is required for enzyme activity (33–35). Complementation of *hem15D* with wild-type human FECH, but not the human FECH variant C406S lacking an essential (33, 36) cluster ligand residue, suggests that the FECH cluster is assembled properly in yeast (Supplementary Fig. S2C).

### Hem15 Is Physically Associated with MICOS Machinery

Putative protein interaction partners of Hem15 were examined by immunoprecipitation (IP) of a FLAG-tagged Hem15 construct. Since attachment of a C-terminal tag to FECH results in an inactive enzyme (37), a version of Hem15 harboring a FLAG epitope tag immediately after the protein’s N-terminal mitochondrial targeting sequence was prepared (Supplementary Fig. S3A). Following in situ proteolytic removal of the targeting sequence after import into the mitochondrion, Hem15 retains the FLAG tag on the structurally disordered amino terminus. This construct, designated Hem15-iFLAG, is stably expressed and functional as judged by its ability to restore heme-independent growth of the *hem15*Δ [*RNR1+*] mutant (Supplementary Fig. S3B and S3C). Clarified mitochondrial lysates from *hem15*Δ [*RNR1+*] cells expressing Hem15-iFLAG were incubated with anti-FLAG affinity resin, and affinity-purified proteins were analyzed by LC-MS/MS to identify co-purifying proteins. In addition to Hem15 and some of its previously known binding partners such as Ppox (Hem14) (17), five out of the eight known subunits of the MICOS complex were identified as well as the IM GTPase Mgm1 (Fig. 3A). In line with these findings, direct co-IP experiments showed the core MICOS subunit Mic60 co-purifies with Hem15-iFLAG (Fig. 3B), further suggesting physical association between Hem15 and MICOS. These findings are consistent with interactions observed in IP experiments with yeast MICOS (38) and human FECH (17, 19).

**Figure 3.**
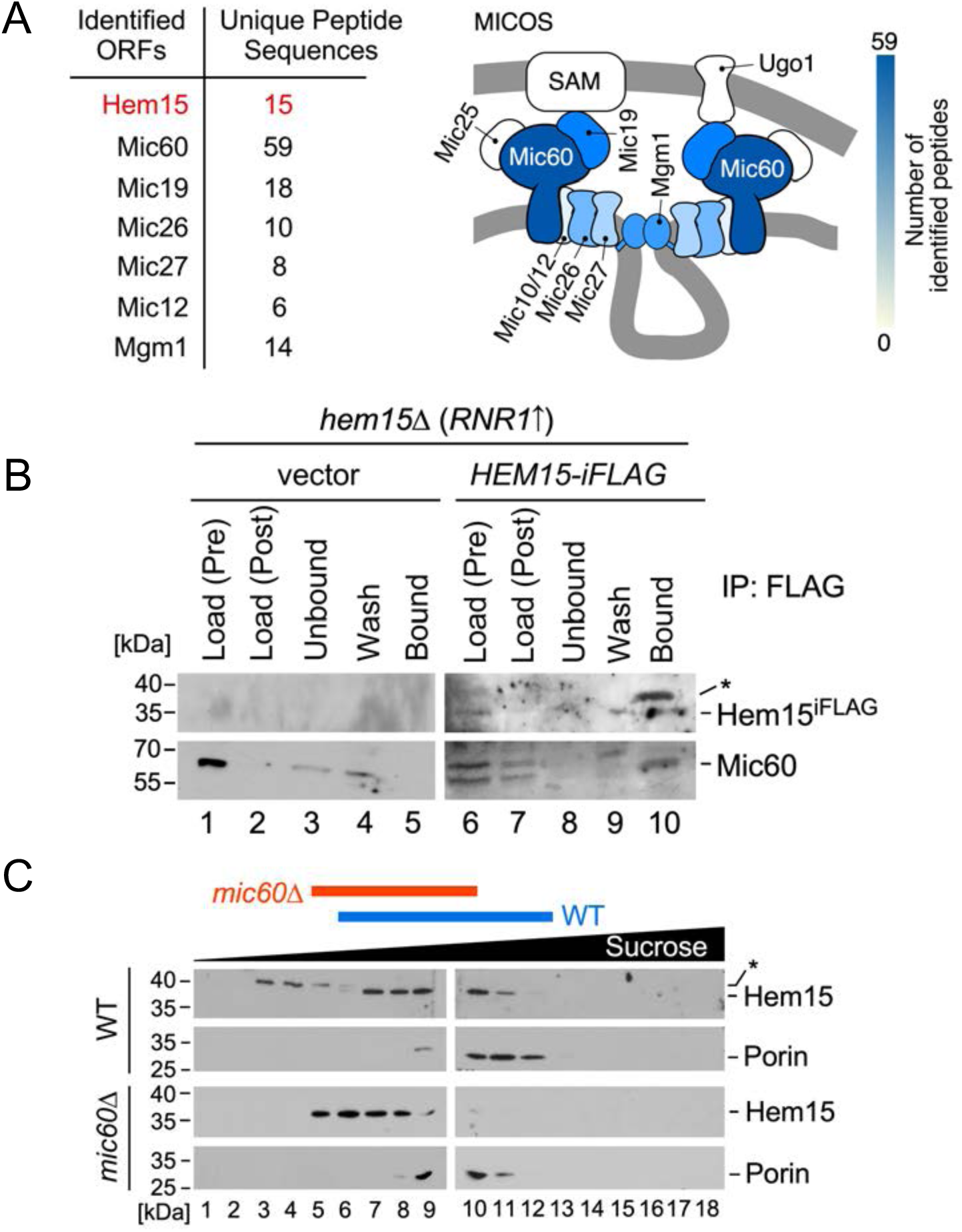
Hem15 physically interacts with MICOS. **(A)** Schematic and table summarizing LC-MS/MS results of immunoprecipitated proteins from mitochondrial lysates of *hem15*Δ cells expressing Hem15-iFLAG. Identified components of the MICOS machinery are color-coded to reflect the abundance of peptides corresponding to each identified subunit. Table shows representative data of two independent biological replicates. **(B)** Co-immunoprecipitation of Hem15-iFLAG and the endogenous Mic60 core subunit of MICOS. Digitonin-solubilized mitochondrial lysates from *hem15*Δ cells expressing Rnr1 and co-expressing either Hem15-iFLAG or empty vector control were incubated with anti-FLAG affinity resin; following the incubation, samples were subjected to SDS-PAGE and analyzed by immunoblotting with indicated antibodies. The blots show 10% of mitochondrial lysate fractions before (Load, Pre) and after (Load, Post) pre-clearance with IgG agarose beads, the unbound fraction after incubation with anti-FLAG affinity resin (Unbound), the whole precipitated fraction from the final wash (Wash), and half of the eluted fraction (Bound). **(C)** Sucrose density gradient ultracentrifugation of digitonin-solubilized Hem15 complexes from mitochondria of WT and *mic60*Δ cells, analyzed as in Fig. 1D. Asterisk marks nonspecific bands.

To examine the functional significance of this Hem15-MICOS association, the impact of *MIC60* deletion on Hem15 was experimentally determined. Steady-state levels of Hem15 were not affected in the *mic60*Δ mutant (Supplementary Fig. S3D). However, the sucrose gradient migration profile of the Hem15 high-mass complex derived from *mic60*Δ mutant mitochondria was altered, indicating a shift toward smaller complex size (Fig. 3C). Notably, this shift in Hem15 oligomer fractionation pattern is similar to that observed for both the H235C Hem15 variant and WT Hem15 in the *hem1*Δ mutant (compare Fig. 1D and 3C). Collectively, these results suggest that Hem15 associates with the MICOS complex and Hem15 oligomerization is impaired in the absence of MICOS.

### MICOS Facilitates Proper Heme Biosynthetic Activity of Hem15

Hem15 overexpression in a *mic60*Δ background results in impaired growth on respiration-forcing carbon sources (Fig. 4A and 4B). This phenotype is independent of the strain, as it was observed for *mic60*Δ strains in both the W303 and BY4741 genetic backgrounds. Both vector-expressing WT and *mic60*Δ cell lysates have similar Fech activity, and overexpression of Hem15 in each cell type results in significantly increased PPIX metalation by Hem15 (Fig. 4C). Strikingly, Hem15-overexpressing *mic60*Δ cells have dramatically lower Fech activity than Hem15-overexpressing WT cells, suggesting an apparent inability to maximize heme production in the absence of Mic60, despite the cells having similar steady-state levels of Hem15 protein (Fig. 4B and 4C). It is unclear whether this apparent inhibition of Hem15 activity in lysates from *mic60*Δ is due to accumulation of a proposed endogenous inhibitor (39), a post-translational modification of Hem15 due to *MIC60* deletion, or another cause.

**Figure 4.**
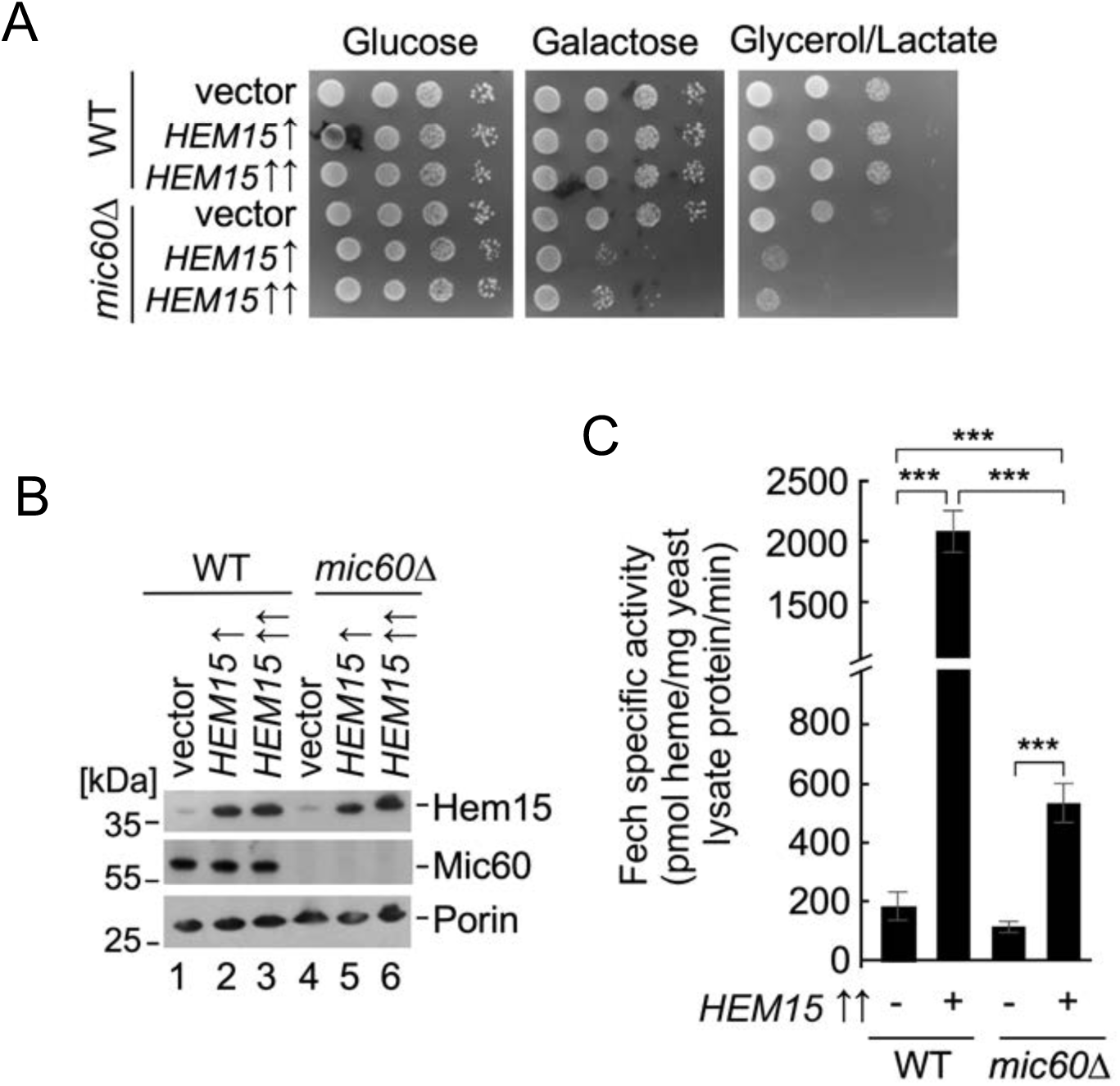
Hem15 activity is impaired in the absence of MICOS. **(A)** Fermentative and respiratory growth of WT and *mic60*Δ cells expressing vector control or Hem15 under control of its native promoter (*HEM15* ↑) or the heterologous *MET25* promoter (*HEM15* ↑↑). Cells were spotted onto SC media and cultured for 2 days (glucose), 3 days (galactose), or 5 days (glycerol/lactate) at 28°C. **(B)** SDS-PAGE immunoblot of indicated proteins in mitochondria from cells described in panel A. **(C)** Ferrochelatase specific activity in mitochondrial lysates from indicated cells. Bars indicate the average and S.D. (error bars) of 3 biological replicates. Asterisks indicate a statistically significant difference by t-test (***p<0.001).

### MICOS Is Required for Optimal Porphyrin Substrate Delivery to Hem15

These data raise questions about the origin of the observed growth defect upon overexpression of Hem15 in MICOS-compromised cells and, more broadly, how the MICOS machinery contributes to Fech activity. Since MICOS plays a key role in maintenance of IM-OM contact sites (21, 22), which are postulated to facilitate bidirectional transport of hydrophobic molecules (e.g., phosphatidic acid and coenzyme Q biosynthetic intermediates) (23, 24), it is possible that heme could also be exported out of mitochondria via the MICOS machinery-facilitated route. In this scenario, impaired heme export due to the absence of MICOS might lead to mitochondrial heme build-up and inhibition of Fech by heme (40), since heme dissociation from FECH is the rate-limiting step in catalysis (41).

To investigate heme homeostasis in WT and MICOS-compromised cells, total steady-state heme levels were measured and found to be similar in WT and *mic60*Δ cells, whether cultured in glucose or galactose media (Supplementary Fig. S4A). To probe heme transport dynamics, the rate of hemylation of a high-affinity genetically encoded heme sensor (HS1) was measured (42, 43). In this assay, heme synthesis is blocked with succinylacetone (SA), an inhibitor of the heme synthetic enzyme Pbgs, and then reinitiated by the inhibitor’s removal from the medium. Upon re-initiation of heme synthesis, the heme occupancy of mitochondrial matrix-and cytosol-targeted HS1 is monitored as a function of time (44). In principle, the rates of heme binding to the sensor reflect the relative rates of heme trafficking from the matrix side of the mitochondrial IM where the active site of Fech is located to the locale of HS1. This assay revealed that hemylation rates of cytosol-and mitochondrial matrix-targeted HS1 in the *mic60*Δ mutant were comparable to those seen in WT cells, indicating that there is no significant dependence of heme export on MICOS (Fig. 5A).

**Figure 5.**
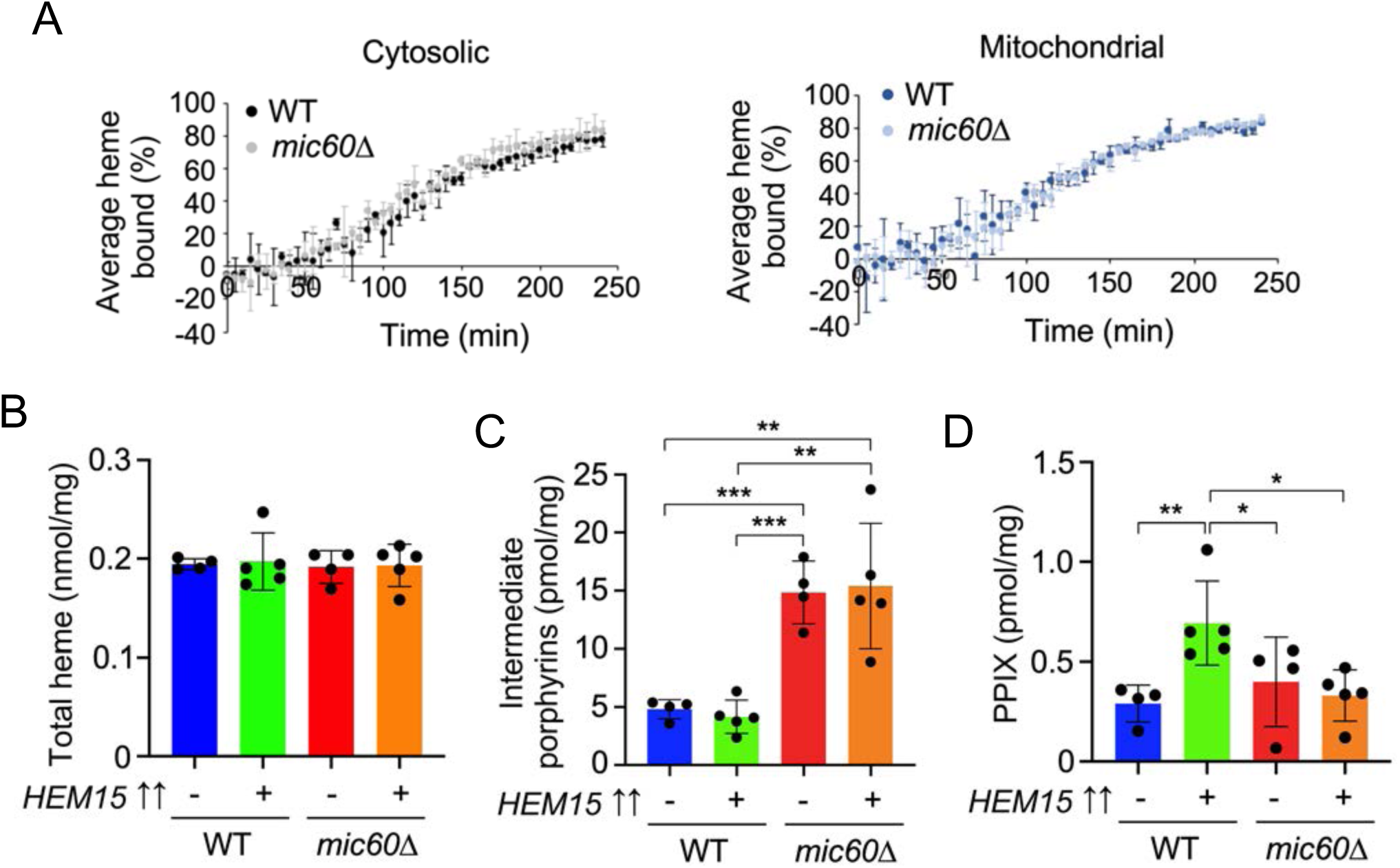
MICOS-deficient cells accumulate reactive porphyrin biosynthetic precursors. **(A)** Relative rates of heme trafficking to the mitochondrial matrix and cytosol in WT and *mic60*Δ cells as measured using the heme sensor HS1. Heme trafficking kinetics data shown represent mean ± S.D. of independent triplicate cultures. **(B-D)** Heme, intermediate (8-, 7-, 6-, 5-, and 4-COOH) porphyrins, and protoporphyrin IX (PPIX) levels in WT and *mic60*Δ cells with and without Hem15 overexpression, analyzed by UPLC. Bars indicate average ± S.D. (error bars) of 4-5 biological replicates measured in technical triplicates. Asterisks indicate a statistically significant difference by t-test (*p<0.05, **p<0.01, ***p<0.001).

When Hem15 is overexpressed in the *mic6*0Δ mutant there is a clear defect in enzymatic activity that is not observed under basal conditions in either WT or *mic6*0Δ cells (Fig. 4C). Yet inter-compartmental heme transport rates between WT and *mic60*Δ cells are nearly equivalent, suggesting that heme distribution kinetics do not appear to be limited by heme synthesis. This observation could indicate that a threshold level of heme must be synthesized prior to its trafficking from Fech.

Another scenario is that the IM disorganization stemming from MICOS loss may negatively impact the function of carrier proteins that transport reactants and intermediates required for heme synthesis, thereby limiting substrate availability to mitochondrial enzymes such as Fech. The iron content measured by ICP-MS was comparable in highly purified mitochondrial fractions from WT and *mic60*Δ cells with or without Hem15 overexpression (Supplementary Fig. S4B), indicating that iron deficiency is not the reason for reduced Fech activity in the Hem15-overexpressing *mic60*Δ mutant. Consistent with this finding, the steady-state levels of the mitochondrial iron transporter Mrs3 were also unchanged in these cells (Supplementary Fig. S4C). Likewise, loss of Mic60 had no appreciable effect on the activity of the iron-sulfur cluster-containing enzyme succinate dehydrogenase, indicating that iron-sulfur cluster biogenesis is not affected (Supplementary Fig. S4D).

To test the availability of the substrate for Fech (PPIX), cellular porphyrin content was analyzed in cells lacking Mic60. Spectroscopy revealed that total porphyrin fluorescence is increased in *mic60*Δ cells compared to WT (Fig. S4E). UPLC was then employed to separate porphyrins to determine which types were causing this elevated fluorescence. While total heme levels measured by UPLC were not significantly different, detailed porphyrin profiling revealed an approximately 3-fold increase in total intermediate porphyrins in *mic60*Δ cells (with or without Hem15 overexpression) when compared to control cells (Fig 5B and 5C) and that the proportions of the 8-, 7-, 6-, 5-, and 4-COOH porphyrins were altered in these cells (Supplementary Fig. S5). Interestingly, PPIX was decreased in *mic60*Δ cells overexpressing Hem15 compared to WT cells overexpressing Hem15 (Fig 5D), suggesting substrate availability may be a factor that limits Fech activity in these cells. These data also support a role for Fech in regulating porphyrinogen homeostasis, as observed in mammalian cells (17).

### MICOS-deficient Cells Expressing Hem15 Have Oxidative Stress, Which Can be Partially Mitigated by a Synthetic Restoration of IM/OM Contact Sites

Heme synthesis pathway intermediates are known to be toxic, in part due to their inherent redox reactivity (5, 45–47), so signs of oxidative damage and stress were evaluated in the absence of Mic60. Hem15-overexpressing *mic60*Δ cells, but not Hem15-overexpressing WT cells, exhibited a significant decrease in aconitase specific activity, consistent with oxidative damage to this superoxide-sensitive metabolic enzyme (Fig. 6A) (while the steady-state levels of aconitase remained unaffected, Fig. S6). Furthermore, *mic60*Δ cells overexpressing Hem15 were significantly less tolerant to acute oxidative insults than WT cells overexpressing Hem15 (Fig. 6B). Consistent with these observations of oxidative stress, supplementation with the antioxidant N-acetylcysteine partially rescued the growth defect of Hem15-overexpressing *mic60*Δ cells on galactose medium (Fig. 6C).

**Figure 6.**
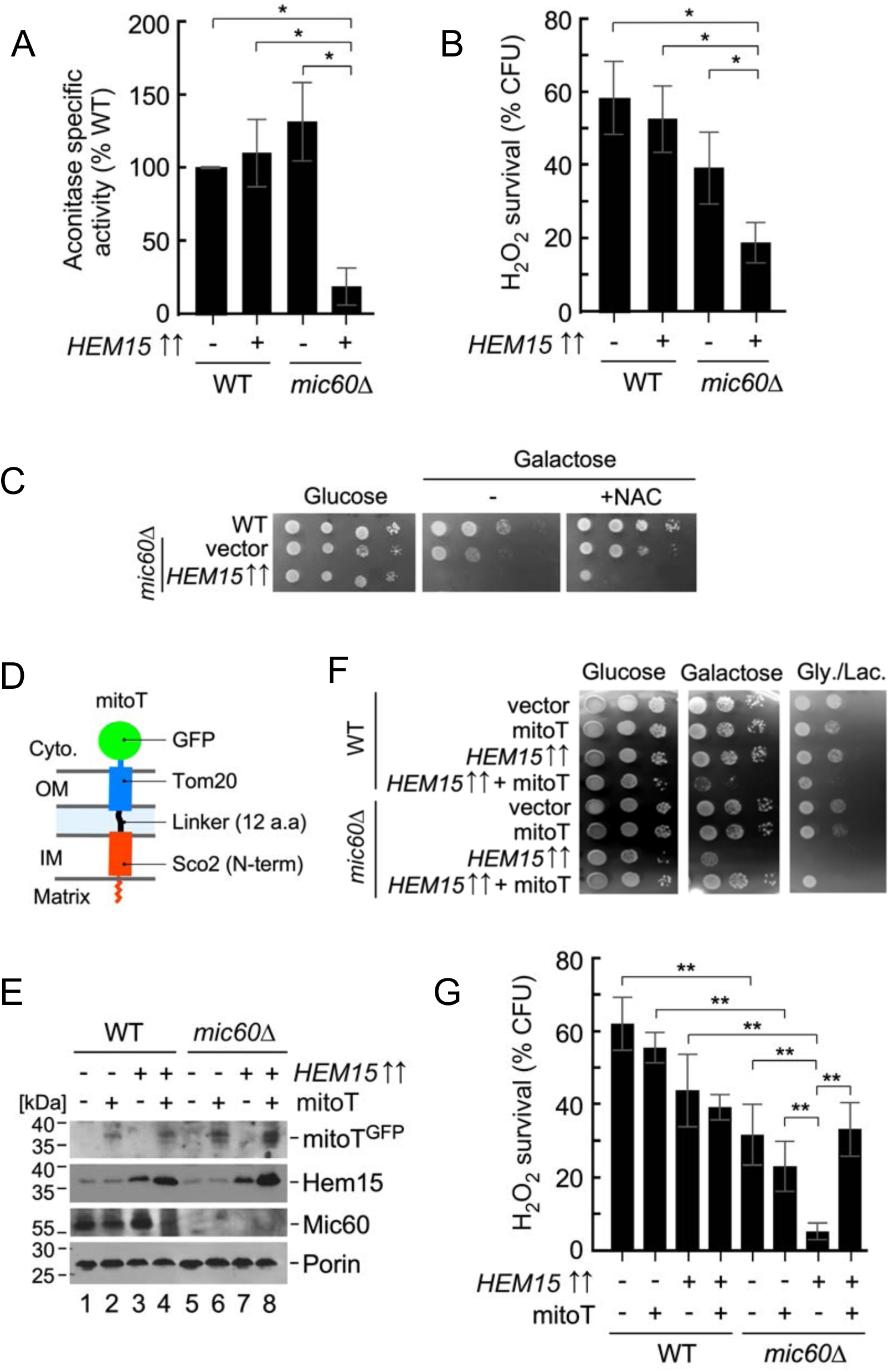
Synthetic intermembrane tether can mitigate oxidative stress in MICOS-deficient cells expressing Hem15. **(A)** Aconitase enzymatic activity in mitochondria isolated from WT and *mic60*Δ cells with and without Hem15 overexpression, harvested after 5 days in culture. Data are mean ± S.D. (n=3 biological replicates). Asterisks indicate a statistically significant difference by one-way ANOVA with Tukey’s post-hoc test (*p<0.05). **(B)** Hydrogen peroxide sensitivity of cells described in panel A. Cells were cultured to mid-log phase, normalized, and acutely treated with 1 mM hydrogen peroxide for 1 h at 28°C. Following treatment, cultures were diluted to 300 cells per sample and plated to assess viable colony forming units after 48 h of growth at 28°C. Bars indicate the average ± S.D. (error bars) of 4 biological replicates. Asterisks indicate a statistically significant difference by one-way ANOVA with Tukey’s posthoc test (*p<0.05). **(C)** Growth test of cells described in panel A with or without the addition of 10 mM N-acetylcysteine (NAC), assessed as in Fig. 4A. **(D)** Schematic depicting mitoT synthetic intermembrane tether. **(E)** SDS-PAGE immunoblot analysis of mitochondria isolated from WT and *mic60*Δ cells expressing vector controls, overexpressing Hem15 or mitoT, or overexpressing both constructs simultaneously. Steady-state levels of indicated proteins were visualized with appropriate antibodies (anti-GFP antibody was used to detect mitoT). The outer mitochondrial membrane protein Porin served as a loading control. **(F)** Growth test of cells described in panel E, assessed as in Fig. 4A. **(G)** Hydrogen peroxide sensitivity of cells described in panel E, handled and analyzed as in panel B.

To test whether restoring intermembrane connectivity in MICOS-deficient cells could mitigate some of these cytotoxic effects, an artificial IM-OM tether, mitoT, was generated (48) (Fig. 6D and 6E). This chimeric protein comprises the N-terminal portion of the IM protein Sco2 (residues 1-112) encompassing the mitochondrial targeting sequence and transmembrane domain, followed by a short (12 amino acid residues) unstructured linker derived from the *E. coli* LacI protein, a transmembrane helix of the OM import receptor protein Tom20 (residues 1-20), and the GFP moiety. The optimal length of the linker region was deduced from the previously reported analogous construct (23). Co-expression of mitoT in Hem15-overexpressing *mic60*Δ cells improved growth on galactose and glycerol-lactate media, reflecting a partial rescue effect (Fig. 6F). Interestingly, mitoT expression had the opposite effect in Hem15-overexpressing WT cells, suggesting that excess intermembrane tethering has a negative effect on the normal function of mitochondria. MitoT expression also resulted in increased steady-state levels of Hem15 protein in both WT and *mic60*Δ cells (Fig. 6E) although the reason for this effect is unclear at present. The rescue appears to be due to a remarkable increase in oxidative stress tolerance in the Hem15-overexpressing *mic60*Δ cells co-expressing mitoT (Fig. 6G).

## DISCUSSION

The heme biosynthetic pathway in metazoans has been extensively studied, including structural characterization of all the enzymes involved (5, 14, 49). However, how heme biosynthetic enzymes and the pathway as a whole are regulated is incompletely understood. In the present study, a series of detailed biochemical and genetic analyses was carried out to better understand the molecular and functional organization of yeast ferrochelatase (Hem15).

Hem15 is shown herein to assemble into a high-mass oligomer. This complex likely represents the functional enzyme, as it is compromised both in *hem1*Δ cells blocked at a very early stage of heme biosynthesis and in cells expressing the catalytically inactive H235C Hem15 mutant. The molecular defect caused by the H235C substitution was also observed (as seen for the human variant H263C (28)), demonstrating the H235 residue is critical for ferrochelatase activity in yeast, likely via proton abstraction from PPIX for metallation and heme formation.

Hem15-deficient cells exhibit a petite-inducing phenotype that can be rescued genetically through overexpression of ribonucleotide reductase subunit Rnr1, a genetic manipulation known to stabilize mitochondrial DNA in petite mutants (32, 50). This finding explains previously described difficulties in establishing a robust genetic complementation of the *hem15D* mutant with plasmid-borne Hem15 under the control of a strong promoter (29). While the exact molecular underpinnings of the petite-inducing phenotype of *hem15D* are currently unclear, they might arise from impaired hemylation of mitochondrial matrix hemoproteins such as the yeast flavohemoglobin Yhb1 or translational activator Mss51 (51, 52). Yhb1 is known to protect cells against oxidative and nitrosative damage, and its function has been linked to the mitochondrial genome (53, 54). Mss51 regulates processing and translation of mRNAs encoding the Cox1 core subunit of respiratory complex IV in a redox-sensitive manner (51, 52), and loss of Mss51 function might negatively impact the mitochondrial genome. Further studies are warranted to test these scenarios.

Another interesting finding from these experiments is that human FECH is able to efficiently replace the yeast enzyme. One significant difference between the yeast and human Fech is the presence of a [2Fe-2S] cluster in the latter enzyme (33–35). This [2Fe-2S] cluster is necessary for enzyme activity and is believed to serve as a sensor for the redox state and/or pH of the mitochondrial matrix (33, 55). Our finding that WT human FECH, but not a mutant bearing the C406S substitution for one of the cluster-binding ligands, complements the yeast deletion mutant suggests that the cluster is successfully formed in the context of yeast mitochondria.

Proteomic analysis suggests the high-mass Hem15 complex contains components of the MICOS machinery. A portion of the core MICOS subunit Mic60 is exposed to the matrix side of the IM (21, 22) and is likely to mediate Hem15-MICOS association. Intriguingly, Mic60 is highly conserved with homologs found in *α*-proteobacteria, wherein the gene clusters with heme biosynthetic genes (56, 57), further indicating a potential functional link between MICOS and the heme biosynthetic pathway. We found that loss of Mic60 affects the high-mass oligomeric species of Hem15 akin to the destabilizing effect of the catalytic H235C Hem15 variant and the Alas (Hem1) deletion mutant. This observation is consistent with the idea that MICOS contributes to the molecular organization of Fech. The yeast MICOS presents as a series of large oligomers (38) and, since Mic60 is about 1.5-fold more abundant than Hem15 (27, 58), it is unlikely that its subunits form stable stoichiometric complexes with Hem15. Instead, our genetic and functional studies suggest a dynamic association between Hem15 and MICOS. In agreement with this notion is the observation that basal Hem15 activity is not significantly affected in the *mic60D* mutant, whereas overexpression of Hem15 causes a profound growth defect in these cells.

Notably, these results are also in agreement with earlier reports suggesting that mammalian FECH is a component of a large complex termed the mitochondrial heme metabolon (17–19). Studies in mammalian cells have shown the heme metabolon contains the first and seventh enzymes of the heme biosynthesis pathway (ALAS and PPOX), succinyl-CoA synthase (SUCLA2), and the porphyrinogen transporter (TMEM14C) (17, 18). In addition, the mitochondrial iron importer mitoferrin (59) as well as two ATP-binding cassette proteins, ABCB7 and ABCB10, (59–61) have also been shown to bind to FECH. Of note, some of these proteins are specific to higher eukaryotes with no apparent orthologs in yeast. Although these reports and the present study cannot be directly compared due to differences in purification strategies (detergents and buffers, expression of epitope-tagged FECH in WT cells versus the deletion mutant, and cutoff stringency), they are consistent in identifying core components of MICOS as associative partners of ferrochelatase, underscoring the conservation and importance of this connection.

These results shed light on the functional significance of the FECH-MICOS association. Studies in iron-sulfur cluster assembly mutants have shown that FECH activity is inhibited by a reversable inhibitor, and the heme synthesis defect and mitochondrial iron accumulation are not correlated (39). Our data suggest that delivery of iron is not affected in MICOS-depleted cells, while PPIX levels are lower than the comparable WT cells. Therefore, we propose a model wherein IM-OM contact sites formed by MICOS facilitate the transfer of intermediate porphyrinogen precursors across the mitochondrial membranes, thereby ensuring optimal substrate delivery to Fech (Fig. 7). While an active role of MICOS in intermediate porphyrinogen transport seems unlikely, the complex may aid the process through bridging the OM and IM to create a proximity conduit between these membranes. Our results showing the rescue of oxidative stress tolerance in Hem15-overexpressing *mic60D* cells co-expressing a synthetic mitoT (48) tether are consistent with this idea. Alternatively, in mammalian cells, MICOS might be required for spatial organization of coproporphyrinogen III and protoporphyrinogen IX transporters in the OM and IM, respectively. Identification of these currently elusive transporters will help to evaluate this possibility. Importantly, the above scenarios are not mutually exclusive. Consistent with these ideas is the finding that MICOS-deficient cells accumulate tetrapyrrole intermediates arising from cellular accumulation of the heme synthesis intermediates during uroporphyrinogen III decarboxylation (Fig. 5C and Supplementary Fig. S5). This situation is similar to what occurs when FECH is overexpressed in cell culture (17). Surprisingly, there is a decrease in PPIX, possibly suggesting less protoporphyrinogen IX production by Cpox. Since sufficient heme is synthesized even in *mic60D* cells under basal conditions, a decrease in Cpox activity could be caused by heme inhibition (62). The MICOS complex has been proposed to play an active role in the transport of phospholipids such as phosphatidic acid (23) and coenzyme Q biosynthetic intermediates (24). Although our data do not exclude the possibility that MICOS is similarly involved in transport of heme biosynthetic intermediates, the observation that mitochondrial heme trafficking rates are normal in the *mic60D* mutant argues against this scenario.

**Figure 7.**
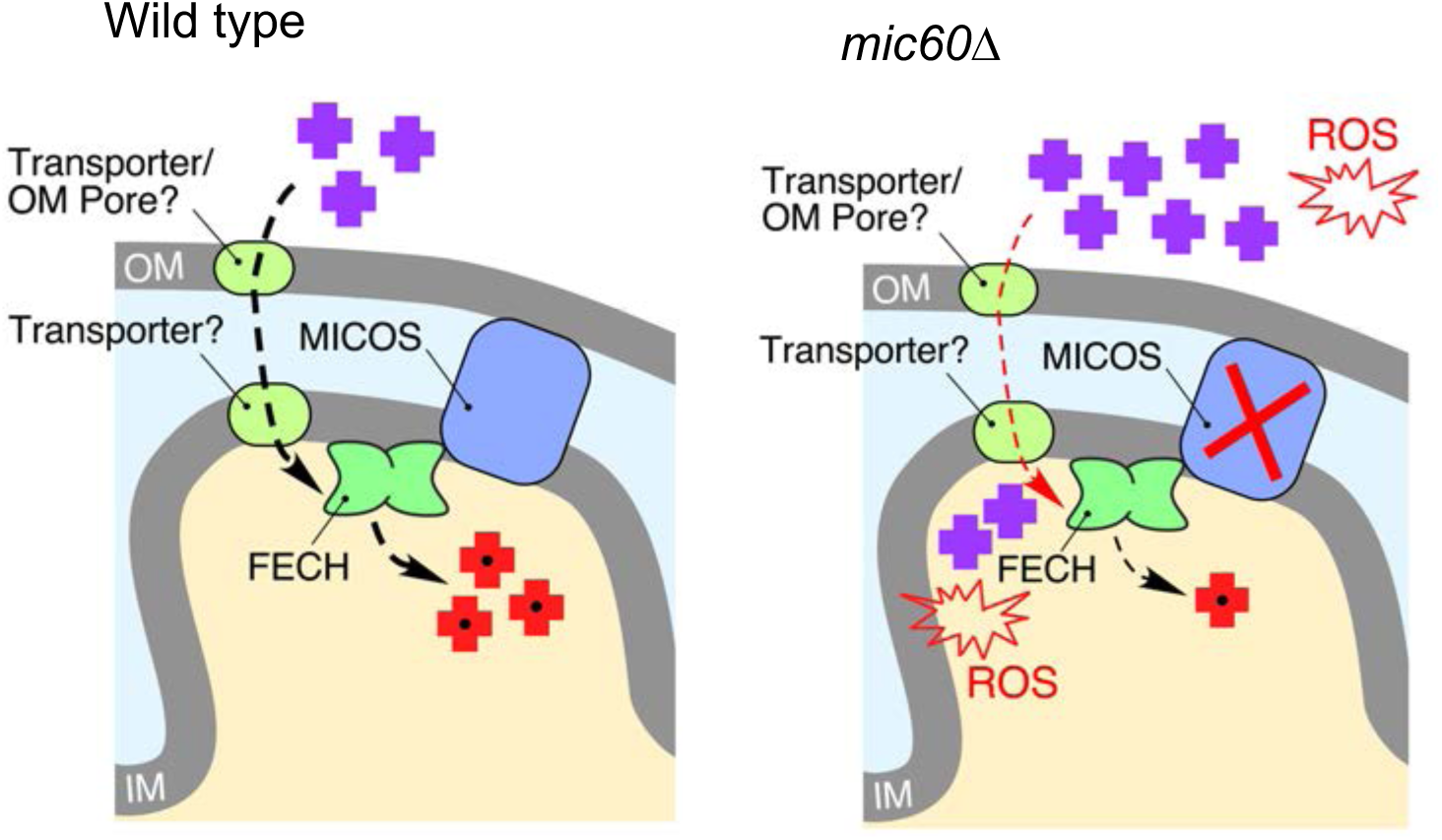
Model for involvement of MICOS in heme biosynthesis. We posit that IM-OM contacts formed by MICOS facilitate the transfer of intermediate porphyrin precursors (purple crosses) across the mitochondrial membranes into the matrix, thus ensuring optimal delivery of substrate to Fech for synthesis of heme (red crosses). Loss of Mic60/MICOS results in accumulation of porphyrin intermediates and subsequent oxidative damage. See Discussion for additional details.

Interestingly, the identification of the conserved dynamin-like GTPase Mgm1/OPA1 as a potential interacting partner of Fech supports earlier proteomic studies in mammalian mitochondria (17, 19). We were unable to biochemically examine the interaction of Mgm1 with Hem15 due to lack of appropriate reagents and enhanced adventitious binding of Mgm1 to various resins; however, our recent study with the HS1 fluorescent heme reporter established that the *mgm1D* mutant exhibits a marked defect in nuclear heme levels (44). These findings further support a model in which IM ultrastructure-related factors like Mgm1 and MICOS may cooperate with Fech to facilitate delivery of its substrate and/or distribution of its product.

## MATERIALS AND METHODS

### Yeast strains, plasmids, and growth conditions

Yeast strains used in this work were of the W303-1B genetic background (*MAT*α *ade2-1 can1-100 his3-11,15 leu2-3 trp1-1 ura3-1*), unless specified otherwise. Plasmids for wild-type and variant Hem15 and human ferrochelatase expression in *S. cerevisiae* were produced as previously described (63, 64). The *HEM15* ORF with its 350-bp native promoter region was subcloned into pRS424 and pRS426 vectors from the pRS316-Hem15 plasmid described before (63). This plasmid also served as a template to generate a pRS426-Hem15 vector in which the *HEM15* ORF is under the control of the heterologous *MET25* promoter.

To generate a plasmid expressing Hem15-iFLAG, the mitochondrial targeting sequence (MTS) of Hem15 (amino acid residues 1-31) with a flanking region containing the FLAG epitope tag at the 3’-end was PCR-amplified from the pRS426-Hem15 plasmid using primers 5’-CGCGGATCCATGCTTTCCAGAACAATCCGTACACAAGGTTCCTTCCTAAGAAGATCACAACTGACCATT-3’ and 3’-CTTGTCGTCATCGTCTTTGTAGTCCTGCATGTTGAATGTAACCGAAAATGATCTTGTAATGGTCAGTTGTGATCTTCT-5’. In a separate reaction, Hem15 excluding the MTS (amino acid residues 31-393) containing the FLAG epitope tag at the 5’-end was PCR amplified from the same plasmid using primers 5’-GACTACAAAGACGATGACGACAAGAATGCACAAAAGAGATCACCCACAGGAATTGTTTTGATGAACATGGGTGGC-3’ and 3’-GCGCTCGAGTCAAGTAGATTCGTGATTGCCAAATACCAATGAAAGGTCCTTTACAGGATCATTGGACTT-5’. Gel-purified products of the reactions described above were fused together by overlap extension PCR using 5’-CGCGGATCCATGCTTTCCAGAACAATCCGTACACAAGGTTCCTTCCTAAGAAGATCACAACTGACCATT-3’ and 3’-GCGCTCGAGTCAAGTAGATTCGTGATTGCCAAATACCAATGAAAGGTCCTTTACAGGATCA TTGGACTT-5’ including 5’-*BamH*I and 3’-*Xho*I restriction sites, respectively. The obtained product was cloned into pRS423 vector under the control of the *MET25* promoter and *CYC1* terminator.

The pRS424-Rnr1 plasmid was generated by subcloning a 2.8-kbp fragment from the YEplac181-Rnr1 vector (32). The pCM185-Mrs3-FLAG plasmid has been described previously (65). All plasmids were validated by DNA sequencing.

Depending on experiment, cells were cultured in yeast extract-peptone (YP) or synthetic complete (SC) media lacking nutrients (amino acids or nucleotides) necessary to maintain plasmid selective pressure (66) containing either 2% glucose, 2% galactose, or 2% glycerol/2% lactic acid mix as the carbon source. To culture heme synthesis-deficient strains, cells were grown in the presence of either hemin or Tween-80/ergosterol/methionine mix as described previously (67, 68). Growth tests to assess respiratory capacity and quantify hydrogen peroxide sensitivity were carried out as before (69, 70), except cells lacking functional Hem15 were cultured in 20 μM hemin. Cultures used for growth tests were grown overnight (or two days in the case of cells lacking functional Hem15) in SC media lacking relevant nutrients to maintain plasmid selection then normalized to OD_600_ of 1 and spotted onto SC plates with or without 20 μM hemin.

### Heme and Porphyrin Analysis

Total cellular heme or protoporphyrin IX were measured using a previously described porphyrin fluorescence assay (44). Briefly, 1×10^8^ log-phase cells were harvested, washed in sterile ultrapure water, resuspended in 500 μL of 20 mM oxalic acid and stored at 4°C overnight (16–18 h) in the dark. The next day, 500 μL of 2 M oxalic acid was added to the cell suspensions. Half the cell suspension was transferred to a heat block set at 95°C and heated for 30 min to demetallate the heme iron to yield fluorescent protoporphyrin IX. The other half of the cell suspension was kept at room temperature for 30 min. All suspensions were centrifuged for 2 min on a table-top microfuge at 21,000 x g and the porphyrin fluorescence spectra (excitation at 400 nm) or emission at 620 nm of 200 μL of each sample was recorded on a Synergy Mx multi-modal plate reader using black Greiner Bio-one flat bottom fluorescence plates. The “boiled” sample provides a measure of total heme and protoporphyrin IX in cells. The “room temperature” sample provides a measure of total protoporphyrin IX in cells. When the “room temperature” sample is subtracted from the “boiled” sample, “total” cellular heme is revealed. Concentrations of heme or protoporphyrin IX are derived from standard curves of known concentrations of heme boiled in oxalic acid similarly to the description above.

For high-resolution heme and porphyrin analysis, yeast cultures (125 mL) were inoculated from a fresh culture at OD_600nm_ of 0.003-0.009 and grown overnight at 37°C and 225 rpm shaking until OD_600nm_ = 1. One hundred mL of culture was harvested by centrifugation at 4000 rpm for 5 min. Cells were then washed in 5 ml ice cold water and re-centrifuged. Pellets were then resuspended on ice in 4 mL lysis buffer (50 mM Tris-MOPS, pH 8.0, 100 mM KCl, 1% sodium cholate) with 40 μL fungal protease inhibitor cocktail (Sigma P-8215) and transferred to 50-mL conical tubes. Glass beads (4 mL, 0.5 mm) were then added, and cells were vortexed for one minute at maximum speed followed by one minute on ice, repeated for a total of 5 vortex cycles. Tubes containing lysate and beads were then centrifuged at ∼9000 x g for 5 min. Supernatant was removed using a 1-mL pipette, taking care not to disturb the pellet. The supernatant was then transferred to 2-ml microcentrifuge tubes and further centrifuged at 15000 x g for 10 min. Supernatant was transferred to fresh microcentrifuge tubes and stored on ice for up to 2 hours during measurements. Total protein concentration of yeast lysate was measured via a NanoDrop spectrophotometer with 1 A = 1 mg/mL setting. Heme and porphyrin analysis were carried out at the University of Utah Center for Iron and Heme Disorders core facility as previously described (11, 17).

### Hem15 activity assays

Lysate volumes ranging from 0-200 μL were brought to a total volume of 850 μL with lysis buffer in 3-mL glass tubes. Master mix (150 μL) consisting of 20 mM ferrous sulfate, ∼1 mM protoporphyrin IX, and 33.3 mM β-mercaptoethanol was added to start the assay. Samples were incubated in the dark at 37°C for 15 min. Heme produced in lysate activity assays was quantified by pyridine hemochromogen assay. To stop the assay reactions and produce the pyridine hemochrome derivative, 1 mL 50% (v/v) pyridine:0.2 N NaOH solution was added to 1-mL assay samples. Heme content was measured via differential spectra of oxidized and reduced samples as described previously (71–73).

### Variant construction, protein expression, purification, and X-ray crystallography

The Hem15 variant H235C was constructed using QuikChange site-directed mutagenesis (Agilent) and verified by sequencing. His-tag mature wild-type and variant Hem15 were produced as previously described (37, 74). In vivo ferrochelatase activity of the variant was assessed by complementation and rescue of a strain of *Escherichia coli* lacking functional ferrochelatase, *ppfC*Δ (75, 76). The H235C variant protein was concentrated to ∼400 μM and crystals were grown by hanging drop with mother liquor composed of 0.1M Bis-Tris pH 6.5, 25% PEG 3350 within 48 hours.

All data sets were collected at the Advanced Photon Source and SER-CAT on beamline 22-ID. Phases were obtained by using a monomer of wild-type Hem15 (PDB ID 1LBQ) as a molecular replacement search model. Molecular replacement was performed using the program CNS (77). Iterative rounds of model building and refinement were performed with the programs COOT (78) and CNS, respectively. Data collection, refinement statistic, and PDB ID for the H235C structure are listed in Table SI. Structural representations were created using PyMol (79).

### Mass spectrometry and data analysis

Affinity purification and mass spectrometry were carried out as described for human ferrochelatase (17), except 1% digitonin was used to solubilize purified mitochondria. Subsequent analysis of candidate hits against the CRAPome database (www.crapome.org) eliminated known contaminants and non-specific interactors.

### Mitochondrial isolation and assays

Mitochondria-enriched fractions were isolated using established protocols (80). For ICP-MS measurements, mitochondrial fractions were further purified using discontinuous 14% / 22% Nicodenz gradients as described (81). Total mitochondrial protein concentrations were determined using the Coomassie Plus kit (Thermo Scientific). Proteins or protein complexes were separated by SDS-PAGE, blue native (BN)-PAGE, or continuous sucrose density gradient ultracentrifugation as previously described (69, 82). Aconitase specific activity was determined as before (83).

### Heme trafficking dynamics assay

Heme trafficking rates were monitored as previously described (44). Briefly, in this three-step assay: 1) heme synthesis is first inhibited with succinylacetone (SA) in sensor-expressing cells, 2) the block in heme synthesis is then removed by resuspending cells into media lacking SA, and 3) the time-dependent change in the heme occupancy of HS1 is monitored. The fractional heme occupancy of the sensor can be determined using previously established sensor calibration protocols (42). The percent of sensor bound to heme (% Bound) is calculated by determining the sensor eGFP/mKATE2 fluorescence ratio (*R*) under a given test condition relative to the eGFP/mKATE2 fluorescence ratio when the sensor is 100% (*R*_max_) or 0% (*R*_min_) bound to heme, as described previously (42–44, 84). *R*_min_ is determined by measuring the HS1 eGFP/mKATE2 ratio in parallel cultures that are conditioned with succinylacetone (SA), which inhibits the second enzyme in the heme biosynthetic pathway, Pbgs (85), and *R*_max_ can be determined by permeabilizing cells and adding an excess of heme to saturate the sensor (42). Given HS1 is quantitatively saturated with heme in the cytosol, nucleus, and mitochondria of WT yeast, *R*_max_ is typically determined by measuring the HS1 eGFP/mKATE2 ratio in parallel WT cultures grown without SA (42).

Growth for the heme trafficking dynamics assay was accomplished by culturing HS1-expressing cells with or without 500 μM SA (Sigma-Aldrich) in SC media lacking leucine. Triplicate 5-mL cultures were seeded at an initial optical density of OD_600nm_ = .01-.02 (∼2-4 x 10^5^ cells/mL) and grown for 14-16 hours at 30°C and shaking at 220 rpm until cells reached a final density of OD_600nm_ ∼ 1.0 (∼2 x 10^7^ cells/mL). After culturing, 1 OD (or ∼2 x 10^7^ cells) were harvested, washed twice with 1 mL of ultrapure water, and resuspended in 1 mL of fresh media. The cells that were pre-cultured without SA provided HS1 *R*_max_ values. The SA-conditioned cells were split into two 500-μL fractions. One fraction was treated with 500 μM SA to give HS1 *R*_min_ values. The other fraction was not treated with SA so that heme synthesis could be re-initiated to give compartment-specific heme trafficking rates. HS1 fluorescence was monitored on 200 uL of a 1 OD/mL (∼2 x 10^7^ cells/mL) cell suspension using black Greiner Bio-one flat bottom fluorescence plates and a Synergy Mx multi-modal plate reader. Fluorescence of eGFP (λ_exc._=488 nm, λ_em._=510 nm) and mKATE2 (λ_exc._=588 nm, λ_em._=620 nm) was recorded every 5 minutes for 4 hours, with the plate being shaken at “medium-strength” for 30 seconds prior to each read. Background fluorescence of cells not expressing the heme sensors was recorded and subtracted from the eGFP and mKATE2 fluorescence values.

### Immunoblotting

Separated proteins were transferred to either nitrocellulose or PVDF membranes, blocked in 5% non-fat milk in PBS with 0.1% Tween-20 and incubated with relevant primary antibodies and goat anti-mouse or goat anti-rabbit horseradish peroxidase-coupled secondary antibodies (Santa Cruz Biotechnology). Proteins of interest were visualized by incubation of membranes with chemiluminescence reagents (Thermo Scientific) and exposure to X-ray film. For assessment of Hem15 endogenous levels, proteins were detected using the Odyssey Fc imaging system (LI-COR Biosciences) and quantified using built-in Image Studio software. The following primary antibodies were used: mouse anti-porin (459500, Thermo Scientific), mouse anti-Cox1 (ab110270, Abcam), mouse anti-Cox2 (ab110271, Abcam), mouse anti-Cox3 (ab110259 Abcam), and mouse anti-FLAG (sc-166355, Santa Cruz Biotechnology). We also used rabbit sera against *S. cerevisiae* Hem15 (produced in the Dailey lab), Rip1 (provided by Dr. D. Winge), Mic60 (provided by Dr. N. Pfanner), Aco1 (provided by Dr. R. Lill), and *β*-subunit of F_1_ ATP synthase (provided by Dr. A. Tzagoloff). All antibodies were tested for reliability to ensure specificity of detection.

## Supporting information

Dietz_Supplemental Information

## ACKNOWLEDGEMENTS

We thank Drs. Dennis Winge, Diane Ward, and John Phillips (University of Utah); Antoni Barrientos (University of Miami); Alexander Tzagoloff (Columbia University); Nikolaus Pfanner (University of Freiburg); and Roland Lill (University of Marburg) for reagents. We also thank Dr. Javier Seravalli and the University of Nebraska-Lincoln Redox Biology Center Biophysics Core for help with ICP-MS analyses and Hector Bergonia at the University of Utah for help with heme and porphyrin analyses.

