## Supplementary material for "Mitochondrial Contact Site and Cristae Organizing System (MICOS) Machinery Supports Heme Biosynthesis by Enabling Optimal Performance of Ferrochelatase": Dietz_Supplemental Information

**Supplemental Table**

**Table S1. Data collection and refinement statistics**

| | |
|---|---|
| *Data Collection* | |
| Beamline | APS 22-ID |
| Space Group | $P2_12_12_1$ |
| Wavelength | 0.98 |
| Resolution Range (Å) | 50.0-2.4 |
| Outer Shell | 2.51-2.4 |
| Unique Observations | 32,665 |
| Completeness (%) | 99.2(96.8)[a] |
| $R_{sym}$ (%)[b] | 6.6(31) |
| CC ½ | 0.99(0.72) |
| Redundancy | 3.5(2.9) |
| $I/\sigma$ | 17.7(2.9) |
| *Refinement* | |
| Unit Cell (*a, b, c*) in Å | 49.9, 51.6, 287.3 ; $\alpha=\gamma=\beta=90^o$ |
| Protein Atoms | 5675 |
| Solvent Atoms | 66 |
| Resolution Limits (Å) | 50.0-2.4 |
| $R_{cryst}$ (%) | 23.2 |
| $R_{free}$ (%) | 29.6 |
| rmsd bonds (Å) | 0.009 |
| rmsd angles ( ° ) | 1.1 |
| average B factor (Å$^2$) | 45 |
| PDB ID code | 7L78 |

[a]Numbers in parentheses denote values for the outermost resolution shell. [b]$R_{sym} = \sum_{hkl}[\sum_I(|I_{hkl,I} - \langle I_{hkl}\rangle|)]/\sum_{hkl,I}\langle I_{hkl}\rangle$, where $I_{hkl}$ is the intensity of an individual measurement of the reflection with indices hkl and $\langle I_{hkl}\rangle$ is the mean intensity of that reflection.

**Supplemental Figures**

**Figure S1, related to Figure 1. Biochemical and structural analysis of Hem15 and its H235C variant. (A)** Indicated amounts of *E. coli*-expressed purified Hem15 along with 100 μg of purified mitochondria from WT cells were analyzed by denaturing gel and immunoblotting with anti-Hem15 antibody. The linear signal intensities were assessed by automated densitometry analysis and used to estimate the relative amount of endogenous Hem15. **(B)** WT mitochondria were solubilized with digitonin, and clarified lysates were fractionated through a 12-50% sucrose gradient by ultracentrifugation. Collected fractions were analyzed by SDS-PAGE and immunoblotting with indicated antibodies. The 440-kDa Porin complex was used for size comparison. **(C)** Ability to rescue *E. coli ppfC*Δ ferrochelatase mutant by yeast Hem15 WT and H235C variant and their ferrochelatase catalytic rate constant. Data are representative of 3 biological replicates. **(D)** Absorbance spectra of *E. coli*-purified yeast Hem15 WT and H235C variant. Dashed line indicates 427 nm Soret band for heme. Spectra are representative of 3 biological replicates. **(E)** Crystal structures (2.4 Å) of Hem15 WT and H235C variant showing views of the active site with the hydrogen-bonding network (dashes) and key residues involved in catalysis (sticks representation).

**Figure S2, related to Figure 2. Rnr1 expression permits complementation of the *hem15*Δ mutant with human FECH. (A)** Heme-dependent fermentative and respiratory growth of WT cells or *hem15*Δ cells expressing vector control or Hem15 (under the control of its native promoter, labeled as *HEM15* ↑), assessed as in Fig. 2C. **(B)** Heme-dependent growth of Rnr1-expressing *hem15*Δ cells harboring yeast or human ferrochelatase, assessed as described in Fig. 2C. **(C)** Heme-dependent growth of WT and untransformed *hem15*Δ cells (-) or *hem15*Δ cells expressing vector control, yeast Hem15 or indicated variants of human FECH, assessed as described in Fig. 1C.

**Figure S3, related to Figure 3. Generation of a functional Hem15-iFLAG construct and steady-state levels of Hem15 in *mic60*Δ cells. (A)** Schematic depiction of the Hem15-iFLAG construct bearing an internal FLAG epitope tag. *MTS*, mitochondrial targeting sequence. **(B)** Heme-dependent glucose growth of WT and *hem15*Δ cells co-overexpressing *RNR1* and either vector control, untagged Hem15, or Hem15-iFLAG under control of the *MET25* promoter, assessed as in Fig. 1C. **(C)** Expression of Hem15-iFLAG in *hem15*Δ cells overexpressing *RNR1* analyzed by SDS-PAGE and immunoblotting with anti-FLAG antibody (and anti-Porin antibody as a loading control). **(D)** Steady-state levels of indicated proteins in mitochondria from WT, *mic60*Δ, and *hem15*Δ cells, visualized by SDS-PAGE and immunoblotting with indicated antibodies.

**Figure S4, related to Figure 5. Analysis of heme, iron, iron-related proteins, and porphyrin fluorescence in MICOS-deficient cells. (A)** Total heme content of WT and *mic60*Δ cells cultured with or without 500 μM succinylacetone (SA) in 2% glucose or 2% galactose. Data are mean ± S.D. of triplicate cultures. *p<0.01 (one-way ANOVA with Bonferroni posthoc test). **(B)** Elemental ICP-MS analysis of iron levels in gradient-purified mitochondria from WT and *mic60*Δ cells expressing vector control or Hem15 under the heterologous *MET25* promoter (*HEM15* ↑↑). Bars indicate the average and S.E.M (error bars) of 3 biological replicates. The right panel shows steady-state levels of indicated proteins in gradient-purified mitochondria (pure mito.) analyzed by SDS-PAGE immunoblot with appropriate antibodies. **(C)** Steady-state levels of indicated proteins in mitochondria from the cells described in panel B analyzed by SDS-PAGE immunoblot with appropriate antibodies to show that Hem15 overexpression increases steady-state levels of mitoferrin protein Mrs3. **(D)** Succinate dehydrogenase specific activity in mitochondrial lysates from cells described in panel B. Bars indicate the average and S.D. (error bars) of 3 biological replicates. **(E)** Fluorescence spectra of steady-state total porphyrinogenes from oxalic acid-digested cell extracts ($1 \times 10^8$ cells/mL) of the indicated cell cultures. The data represent mean ± S.D. of independent triplicate cultures.

**Figure S5, related to Figure 5. Detailed analysis of identifiable porphyrin biosynthetic precursors in MICOS-deficient cells.** Indicated intermediate (8-, 7-, 6-, 5-, and 4-COOH) porphyrins, as derived from pathway intermediate porphyrinogens, were analyzed by UPLC. Bars indicate average ± S.D. (error bars) of 4-5 biological replicates measured in technical triplicates. Asterisks indicate a statistically significant difference by t-test (*$p<0.05$, **$p<0.01$).

**Figure S6, related to Figure 6. Steady-state aconitase levels are unaffected in Hem15-overexpressing *mic60*Δ cells.** Steady-state levels of indicated proteins in mitochondria from WT and *mic60*Δ cells expressing vector control or Hem15 under control of its native promoter (*HEM15* ↑) or the heterologous *MET25* promoter (*HEM15* ↑↑), analyzed by SDS-PAGE immunoblot with appropriate antibodies. Asterisk indicates non-specific cross-reacting bands.

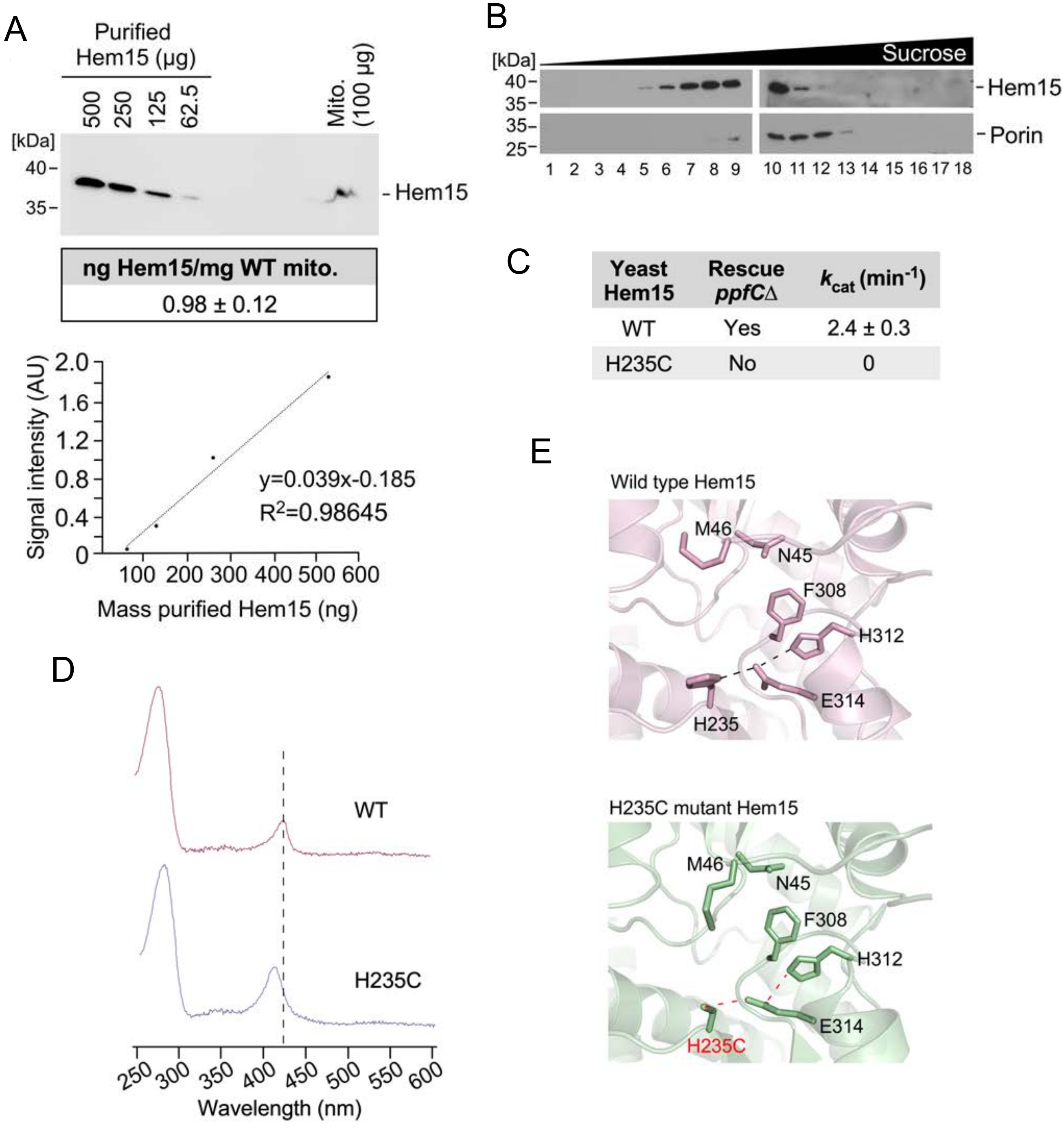
A
Purified Hem15 (μg)
500
250
125
62.5
Mito. (100 μg)
[kDa]
40
35
Hem15
ng Hem15/mg WT mito.
0.98 ± 0.12
Signal intensity (AU)
2.0
1.6
1.2
0.8
0.4
0
100 200 300 400 500 600
Mass purified Hem15 (ng)
y=0.039x-0.185
R2=0.98645
B
[kDa]
Sucrose
40
35
35
25
Hem15
Porin
1 2 3 4 5 6 7 8 9 10 11 12 13 14 15 16 17 18
C
Yeast Hem15
Rescue ppfCΔ
kcat (min-1)
WT
Yes
2.4 ± 0.3
H235C
No
0
D
WT
H235C
250 300 350 400 450 500 550 600
Wavelength (nm)
E
Wild type Hem15
M46
N45
F308
H312
E314
H235
H235C mutant Hem15
M46
N45
F308
H312
E314
H235C

Figure S1

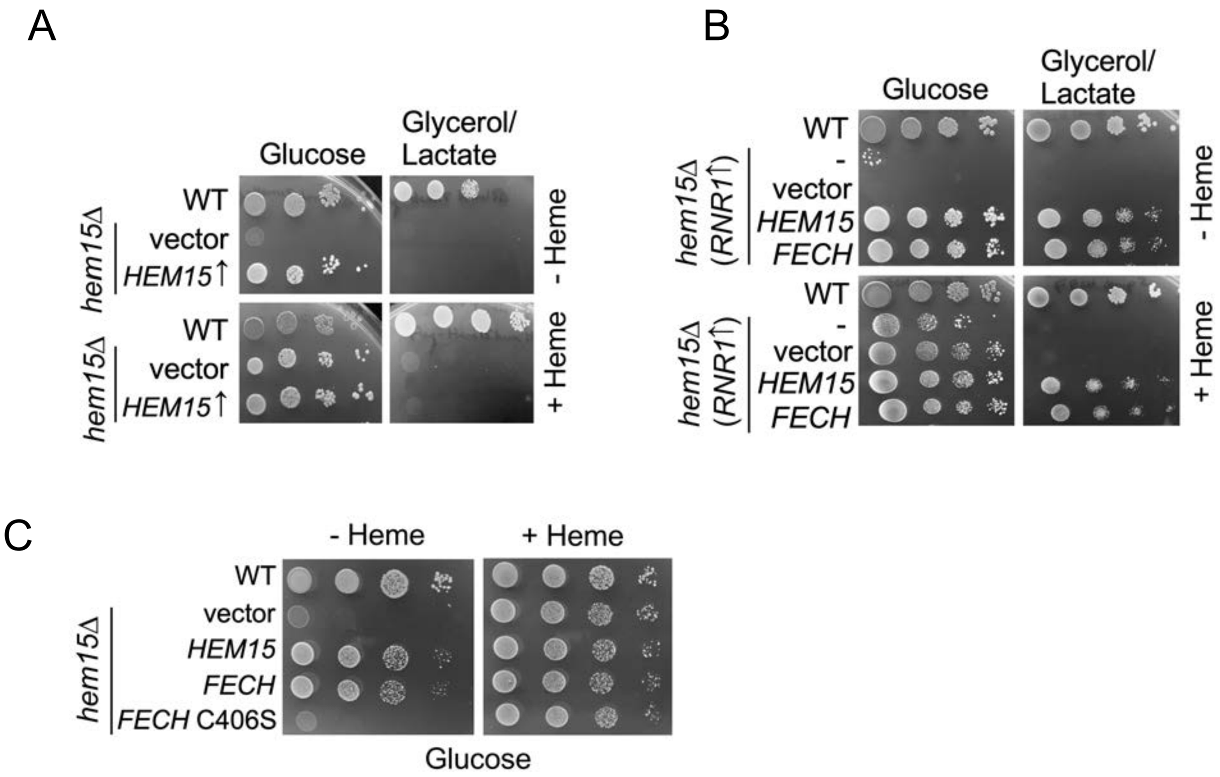
A
Glucose
Glycerol/
Lactate
WT
hem15Δ
vector
HEM15↑
- Heme
WT
hem15Δ
vector
HEM15↑
+ Heme
B
Glucose
Glycerol/
Lactate
WT
hem15Δ
(RNR1↑)
-
vector
HEM15
FECH
- Heme
WT
hem15Δ
(RNR1↑)
-
vector
HEM15
FECH
+ Heme
C
- Heme
+ Heme
WT
hem15Δ
vector
HEM15
FECH
FECH C406S
Glucose

Figure S2

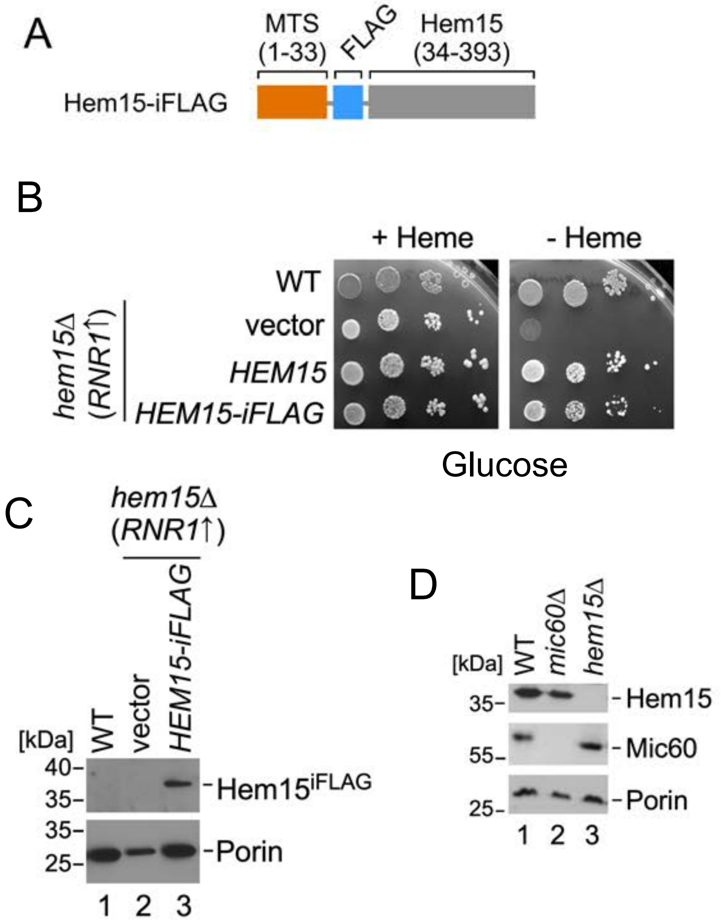
A
MTS
(1-33)
FLAG
Hem15
(34-393)
Hem15-iFLAG
B
+ Heme
- Heme
WT
vector
HEM15
HEM15-iFLAG
hem15Δ
(RNR1↑)
Glucose
C
hem15Δ
(RNR1↑)
[kDa]
WT
vector
HEM15-iFLAG
40-
35-
35-
25-
Hem15iFLAG
Porin
1 2 3
D
[kDa]
WT
mic60Δ
hem15Δ
35-
55-
25-
Hem15
Mic60
Porin
1 2 3

Figure S3

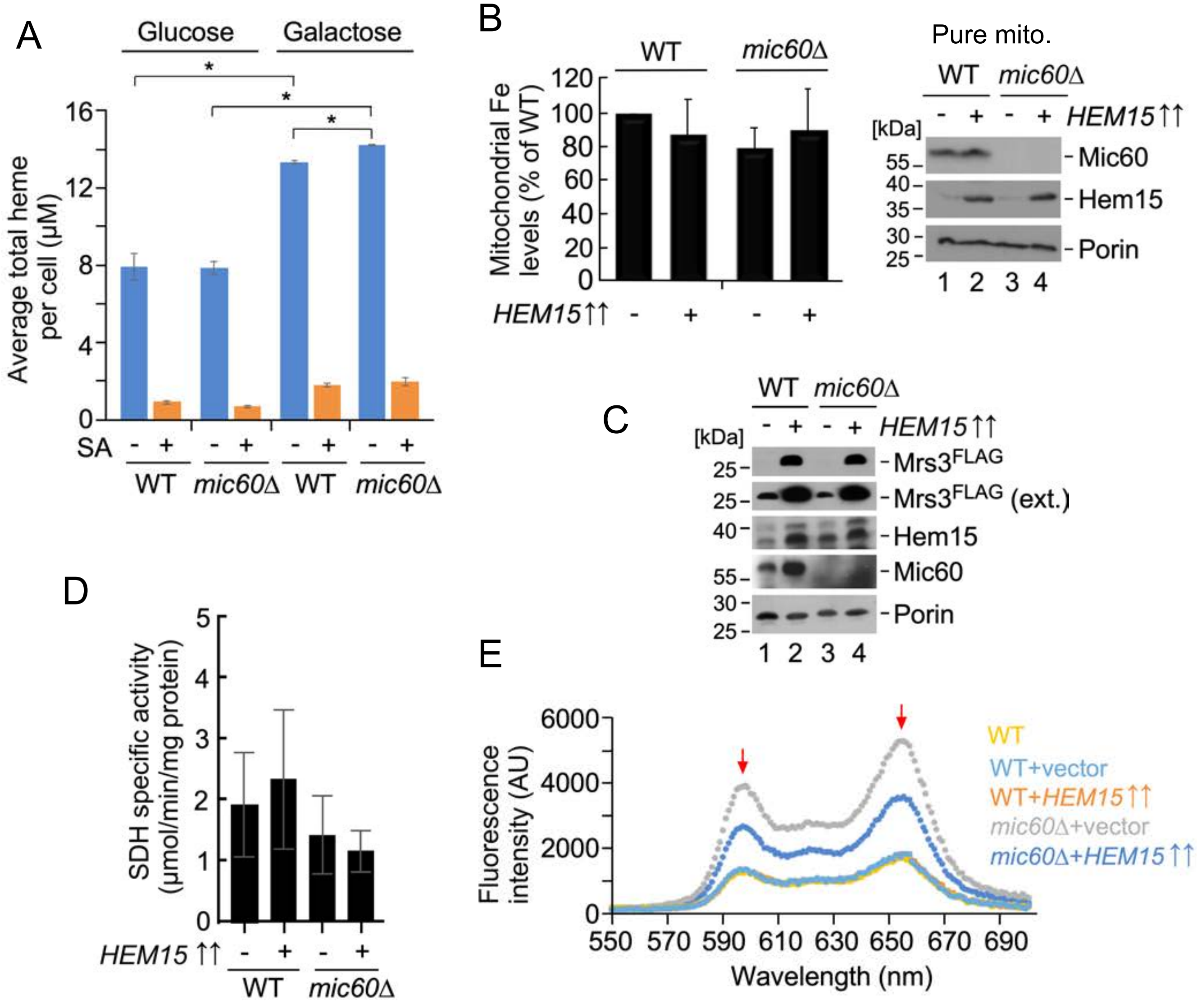
A
Glucose
Galactose
*
*
*
16
12
8
4
0
Average total heme per cell (µM)
SA - + - + - + - +
WT mic60Δ WT mic60Δ
B
WT
mic60Δ
120
100
80
60
40
20
0
Mitochondrial Fe levels (% of WT)
HEM15↑↑ - + - +
Pure mito.
WT mic60Δ
- + - + HEM15↑↑
[kDa]
55
40
35
30
25
Mic60
Hem15
Porin
1 2 3 4
C
WT mic60Δ
- + - + HEM15↑↑
[kDa]
25
25
40
55
30
25
Mrs3FLAG
Mrs3FLAG (ext.)
Hem15
Mic60
Porin
1 2 3 4
D
5
4
3
2
1
0
SDH specific activity (µmol/min/mg protein)
HEM15 ↑↑ - + - +
WT mic60Δ
E
6000
4000
2000
0
Fluorescence intensity (AU)
550 570 590 610 630 650 670 690
Wavelength (nm)
WT
WT+vector
WT+HEM15↑↑
mic60Δ+vector
mic60Δ+HEM15↑↑

Figure S4

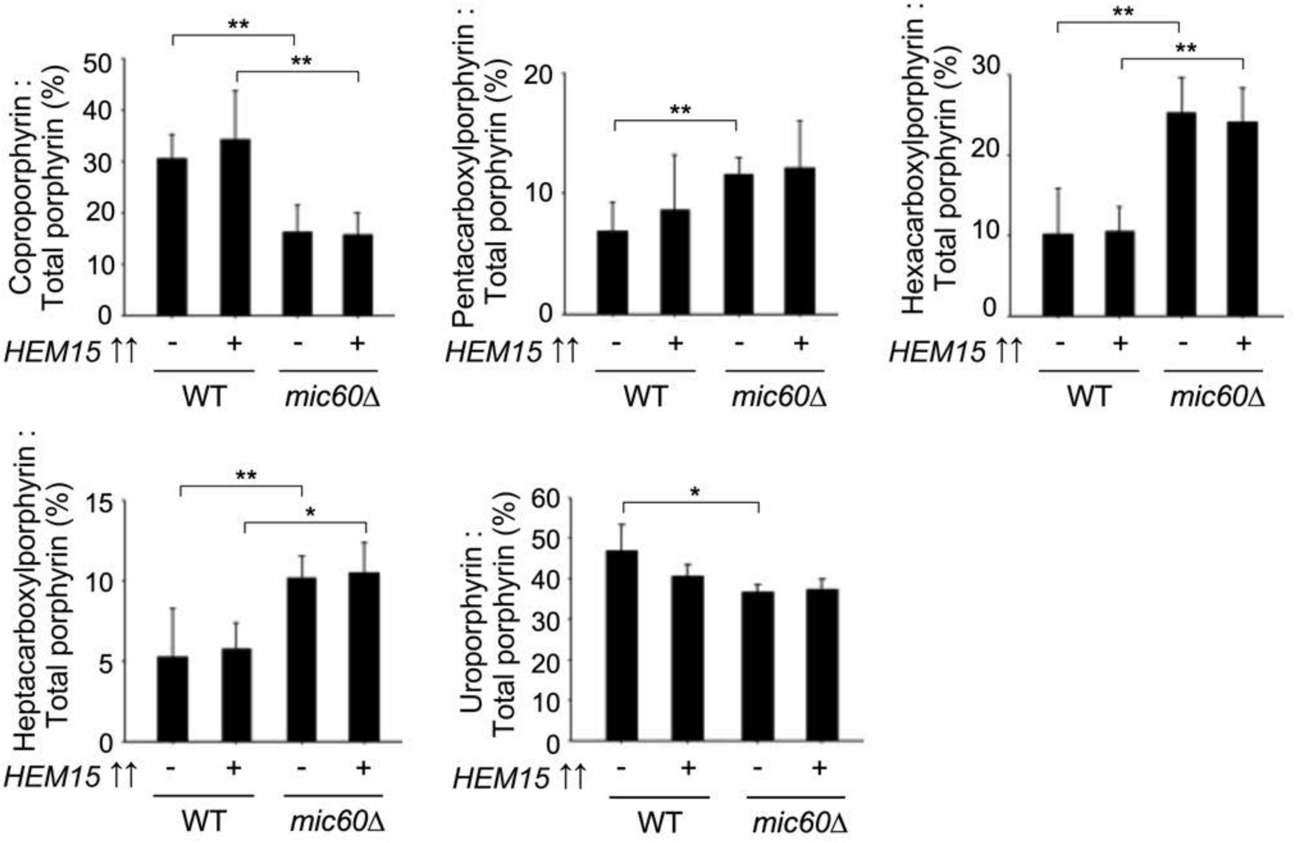
Coproporphyrin : Total porphyrin (%)
Pentacarboxylporphyrin : Total porphyrin (%)
Hexacarboxylporphyrin : Total porphyrin (%)
Heptacarboxylporphyrin : Total porphyrin (%)
Uroporphyrin : Total porphyrin (%)
HEM15 ↑↑
WT
mic60Δ

Figure S5

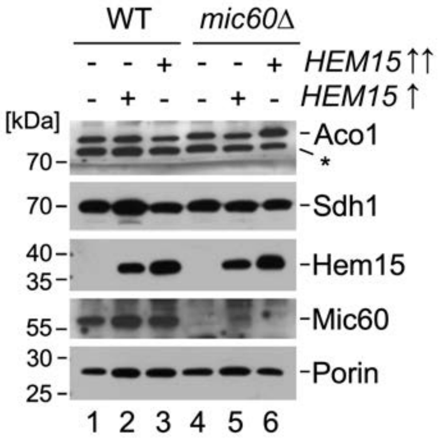

WT
mic60Δ
- - + - - + HEM15↑↑
- + - - + - HEM15↑
[kDa]
Aco1
*
70
Sdh1
70
40
35
Hem15
55
Mic60
30
25
Porin
1 2 3 4 5 6


Figure S6
